# E1: Retrieval-Augmented Protein Encoder Models

**DOI:** 10.1101/2025.11.12.688125

**Authors:** Sarthak Jain, Joel Beazer, Jeffrey A. Ruffolo, Aadyot Bhatnagar, Ali Madani

## Abstract

Large language models trained on natural proteins learn powerful representations of protein sequences that are useful for downstream understanding and prediction tasks. Because they are only exposed to individual protein sequences during pretraining without any additional contextual information, conventional protein language models suffer from parameter inefficiencies in learning, baked-in phylogenetic biases, and functional performance issues at larger scales. To address these challenges, we have built Profluent-E1, a family of retrieval-augmented protein language models that explicitly condition on homologous sequences. By integrating retrieved evolutionary context through block-causal multi-sequence attention, E1 captures both general and family-specific constraints without fine-tuning. We train E1 models on four trillion tokens from the Profluent Protein Atlas and achieve state-of-the-art performance across zero-shot fitness and unsupervised contact-map prediction benchmarks – surpassing alternative sequence-only models. Performance scales with model size from 150M to 600M parameters, and E1 can be used flexibly in single-sequence or retrieval-augmented inference mode for fitness prediction, variant ranking, and embeddings for structural tasks. To encourage open science and further development in retrieval-augmented protein language models, we release three models for free research and commercial use at https://github.com/Profluent-AI/E1.

## Introduction

Proteins are fundamental components of the molecular machinery of life, driving biological processes such as molecular transport, enzyme catalysis, immune response, and gene regulation. Their diverse functions underpin applications across many industries – from pharmaceuticals to agriculture –enabling gene therapies, vaccines, and industrial enzymes. To harness these functions, protein engineers design, modify, or select amino acid sequences that fold into proteins with desired activities. However, mapping sequence to function remains a major challenge, and many traditional engineering strategies still rely on random mutagenesis and high-throughput screening to identify suitable candidates.

Protein language models (PLMs) offer a data-driven framework for modeling the relationships between protein sequence, structure, and function. Trained in a self-supervised manner on large databases of natural protein sequences, PLMs learn evolutionary patterns shaped by natural selection over billions of years [1]. In particular, single-sequence models like ESM-2/C [2–6] trained with masked language modeling objectives produce likelihoods that correlate well with function and internal representations that capture sequence–structure and sequence–function relationships. These models have shown strong performance in protein engineering tasks such as ranking variants by fitness and predicting structure [3, 7].

Despite their success, single-sequence PLMs face key limitations. Firstly, all evolutionary context must be compressed into model parameters, so protein families that are underrepresented in the pre-training data are poorly captured. Fine-tuning can improve performance; however, it is computationally costly, can erase more general protein knowledge, and is infeasible for data-limited families [8]. Second, PLMs model the data distribution itself, meaning they can reflect biases from phylogeny, genetic drift, or sampling, rather than functional constraints [9–11].

To overcome these limitations, recent approaches have incorporated explicit evolutionary context through retrieval augmentation. Retrieval-augmented PLMs (RA-PLMs) enhance standard single-sequence models by providing homologous sequences during training and inference. This allows the model to leverage evolutionary context directly. Notable examples include the MSA Transformer [12], which uses multiple sequence alignments as context, and PoET [13], which employs alignment-free concatenations of homologous sequences. By conditioning on retrieved sequences, RA-PLMs address these challenges:

- **Encoding evolutionary information**. Retrieved homologs provide direct evolutionary context, enabling RA-PLMs to represent both broad and family-specific patterns without overfitting.
- **Contextualizing family-specific fitness**. The model can situate a query sequence within its family’s landscape at inference time, avoiding costly family-specific fine-tuning and supporting low-data applications.
- **Reducing sampling bias**. Conditioning on multiple homologs emphasizes functionally relevant coevolutionary signals while diminishing non-selective sampling or phylogenetic noise.

Empirically, RA methods have proven to be highly effective: multi-sequence attention underlies AlphaFold2’s [14] state-of-the-art structure prediction, and retrieval strategies have shown strong performance in PLMs directly [12, 13, 15, 16]. Beyond predictive accuracy, RA models offer practical flexibility – a single pretrained model can specialize dynamically for specific families or tasks, capturing deep coevolutionary relationships without further training.

In this work, we introduce `Profluent-E1`, a new family of retrieval-augmented protein encoder models trained with a masked language modeling objective. We leverage Profluent’s large-scale Protein Atlas [11] and introduce targeted architectural and training innovations that yield more performant RA-PLMs. `E1` achieves state-of-the-art performance among models trained exclusively on sequence data. On the Protein Gym benchmark for zero-shot fitness prediction, `E1` models outperform the ESM family [2, 3] in single-sequence mode and surpass other retrieval-based models, including PoET [13] and MSA Pairformer [15], when augmented with homologs. `E1` also achieves superior performance in unsupervised contact-map prediction, again outperforming the ESM family in single-sequence mode and showing substantial additional gains with retrieval. We also observe that the performance of our models scales with the number of parameters. We release three `E1` variants – 150M, 300M, and 600M parameters – freely for research and commercial use, enabling immediate application to tasks such as fitness prediction, structure prediction, and representation learning.

## Model

### Architecture

`E1` is a family of retrieval-augmented protein encoder models trained with bidirectional attention and a masked language modeling objective. In contrast to standard protein encoder models like ESM-2 [3], these models leverage sequence homologs as part of their inference context to generate better representations for a given sequence of interest, allowing for in-context learning. Note that we do not require the homologous sequences to be aligned with each other, in contrast to models like MSA Transformer [12] and MSA Pairformer [15]. To test whether the model performance scales with number of parameters, we trained three different sizes of `E1` models: 150M, 300M, and 600M parameters.

The model takes as input a sequence of protein sequences (for example, MLFH,MIIVR,MFHK) with each individual sequence wrapped in special tokens (`<bos>1MLFH2<eos><bos>1MIIVR2<eos><bos>1MFHK2<eos>`) to mark the start and end of the sequence. Embeddings of these tokens are then passed to the model. Each token in the same protein sequence also shares a sequence ID, which is then embedded and supplied to the model to distinguish between different protein sequences within a multi-sequence instance. We allow up to 512 individual sequences within a single multi-sequence instance. `E1` model family is implemented using a standard Transformer-based architecture [17, 18], augmented with a block causal attention mechanism that enables residues in different homologous sequences to attend to one another. For efficiency, this global attention is not applied in every layer. Instead, we adopt an alternating attention architecture [19]: global block-causal attention is used every three layers, while all other layers use intra-sequence attention, where residues attend only to other residues within the same protein sequence.

We use standard Rotary Position Embedding (RoPE) [20] to encode positional information. For layers using intra-sequence attention, each protein sequence restarts position IDs at one, whereas for global-attention layers, the position ID corresponds to the absolute position of the token within the full concatenated multi-sequence input.

### Training

The `E1` family of models was trained using a standard masked language modeling objective [18], in which input tokens are randomly selected and replaced with noisy variations. A language modeling head (a single hidden layer MLP) is then applied on top of the final-layer token representations to predict the probability of the true amino acid at each selected position. During training, we linearly decreased the noise fraction (the fraction of tokens replaced in the input) from 25% to 15% for the first 250 billion tokens; after that, it remained fixed at 15%. We followed the standard BERT masking policy: 80% of selected tokens were replaced with a special mask token, 10% were replaced with a random amino acid, and the remaining 10% were left unchanged. All three `E1` models were trained for 4 trillion tokens (batch size = 2^20^ tokens) using a warmup-stable-decay learning rate schedule [21] and Stable AdamW optimizer [22], on clusters of H100/H200 GPUs – for example, `E1` 600M was trained on a cluster of 64 H100s for 25 days.

### Training Data Construction

To construct multi-sequence instances for training, we adopt the strategy introduced by the PoET model [13]. We used sets of homologous sequences derived from the PPA-1 [11] and UniRef Version 2411 [23] datasets. Both PPA-1 and UniRef are clustered at multiple sequence identity thresholds, including at 50% and 90% identity. For each 50% ID cluster representative, we search it against all other 50% ID cluster representatives in the respective datasets using Diamond [24], returning a set of possible homologs. To construct a training instance, we first randomly sample one of these homolog sets (with probability inversely proportional to the size of the set) and then replace each 50% ID cluster representative with a randomly picked sequence from the associated 50% ID sequence cluster (weighted inversely by the size of its 90% ID subcluster). Finally, we subset the resulting sequences to ensure that the concatenated multi-sequence instance remains within a prescribed length budget.

We employed a curriculum learning strategy where we gradually increase the total length and number of sequences in a multi-sequence instance: from 8192 to 32768 and from 2 to 512 respectively. This enabled the model to achieve state of the art performance in both single sequence mode (where no homologous sequences are passed during inference) and retrieval-augmented mode. During training, we exclusively trained on instances from PPA-1 for the first 1.5 trillion tokens. Thereafter, we mixed in instances from UniRef in a 60:40 ratio for the remainder of the training duration.

## Results

### A. E1 models enable state of the art zero-shot substitution effect prediction

Protein language models have been shown to be effective zero-shot fitness predictors for local mutational landscapes. In addition, prior work [12, 13, 16, 25–27] has shown that addition of evolutionarily related sequences (either unaligned or in the form of an MSA) during inference can improve the model’s performance. In this section, we use the 217 Deep Mutational Scan substitution assays from the ProteinGym (v1.3) benchmark [28] to evaluate the performance of `E1` models in both single-sequence and retrieval-augmented modes. We use the masked marginal method [7] to compute scores for each variant of the wildtype protein sequence and evaluate performance using Spearman correlation and the normalized discounted cumulative gain (NDCG) metric against ground truth fitness values. The latter metric measures the ability of the model to rank high fitness sequences first and is more practically relevant for protein design tasks.

#### Sampling homologs for inference

For evaluation in retrieval-augmented mode, we follow the PoET strategy [13] and prepend the masked variants of the wildtype sequence with homologous sequences sampled from ColabFold derived MSAs [29] constructed using Uniref100 v2104. Homologs are sampled with weights inversely proportional to the number of their neighbors (sequences in the MSA that are at least 80% identical to them)and are additionally constrained to satisfy a specified maximum similarity to the wildtype sequence.. We ensemble 15 prompts corresponding to 3 different total-token-length budgets and 5 different maximum query-similarity thresholds ({6144, 12288, 24576} × {1.0, 0.95, 0.9, 0.7, 0.5}).

#### Results

In Table 1, we observe that `E1` models outperform all ESM-2 and ESMC family models in single-sequence mode at comparable model sizes, indicating that `E1` can be used as a drop-in replacement for existing single-sequence encoder models without loss of performance. When evaluated with homologs at inference time, the `E1` models substantially outperform corresponding single-sequence metrics and achieve state of the art performance relative to similar publicly available models, i.e., models that only take homologous sequences as additional context during inference, like MSA Pairformer and PoET^∗^. In Table 2, we further observe that switching from single-sequence to retrieval-augmented mode yields consistent improvements for assays with low and medium MSA depth. On average, the larger `E1` models also tend to perform better, indicating continued benefits of scaling up retrieval-augmented PLMs.

**Table 1.** Average Spearman correlation and NDCG@10 between model-predicted scores and Protein Gym experimental fitness values.

| Model Name | Model Inputs | Spearman Correlation |  |  |  |  |  | NDCG@10<br>Average |
| --- | --- | --- | --- | --- | --- | --- | --- | --- |
|  |  | Average | Activity | Binding | Expression | Organismal<br>Fitness | Stability |  |
| Inference with query sequence only |  |  |  |  |  |  |  |  |
| ESM2-150M | Sequence Only | 0.387 | 0.391 | 0.326 | 0.402 | 0.305 | 0.51 | 0.729 |
| ESM2-650M | Sequence Only | 0.414 | 0.425 | **0.337** | 0.415 | 0.368 | 0.523 | 0.747 |
| ESM2-3B | Sequence Only | 0.406 | 0.417 | 0.321 | 0.403 | **0.378** | 0.509 | **0.755** |
| ESMC-300M | Sequence Only | 0.406 | 0.423 | 0.315 | 0.408 | 0.36 | 0.526 | 0.746 |
| ESMC-600M | Sequence Only | 0.405 | 0.423 | 0.294 | 0.42 | 0.362 | 0.528 | 0.746 |
| E1 150M | Sequence Only | 0.401 | 0.426 | 0.325 | 0.420 | 0.304 | 0.532 | 0.744 |
| E1 300M | Sequence Only | 0.416 | **0.438** | 0.332 | 0.430 | 0.346 | 0.537 | 0.748 |
| E1 600M | Sequence Only | **0.420** | 0.415 | 0.330 | **0.441** | 0.366 | **0.548** | 0.749 |
| Inference with Homologous Sequences / MSA in-context |  |  |  |  |  |  |  |  |
| MSA Pairformer | Sequence + MSA | 0.45 | 0.49 | 0.35 | 0.44 | 0.46 | 0.51 | — |
| PoET | Sequence + Homologs | 0.470 | 0.494 | 0.396 | 0.466 | 0.475 | 0.519 | 0.784 |
| E1 150M | Sequence + Homologs | 0.473 | 0.498 | 0.408 | 0.468 | 0.477 | 0.514 | 0.785 |
| E1 300M | Sequence + Homologs | 0.475 | **0.501** | **0.410** | 0.468 | 0.474 | 0.523 | 0.787 |
| E1 600M | Sequence + Homologs | **0.477** | **0.501** | 0.404 | **0.469** | **0.478** | **0.532** | **0.788** |

**Table 2.** Average Spearman correlation between model-predicted scores and Protein Gym experimental fitness values broken down by Taxon and MSA Depth.

| Model Name | Model Inputs | Spearman Correlation by Taxon |  |  |  | Spearman Correlation by MSA Depth |  |  |
| --- | --- | --- | --- | --- | --- | --- | --- | --- |
|  |  | Human | Other Eukaryote | Prokaryote | Virus | Low | Medium | High |
| Inference with query sequence only |  |  |  |  |  |  |  |  |
| ESM2-150M | Sequence Only | 0.45 | 0.475 | 0.398 | 0.157 | 0.319 | 0.359 | 0.494 |
| ESM2-650M | Sequence Only | 0.457 | 0.486 | 0.458 | 0.261 | 0.338 | 0.409 | 0.513 |
| ESM2-3B | Sequence Only | 0.442 | 0.477 | 0.458 | 0.294 | 0.336 | 0.423 | 0.485 |
| ESMC-300M | Sequence Only | 0.468 | 0.481 | 0.441 | 0.242 | 0.337 | 0.399 | 0.520 |
| ESMC-600M | Sequence Only | 0.462 | 0.481 | 0.459 | 0.241 | 0.331 | 0.407 | 0.515 |
| E1 150M | Sequence Only | 0.455 | 0.515 | 0.413 | 0.188 | 0.342 | 0.373 | 0.514 |
| E1 300M | Sequence Only | 0.466 | 0.513 | 0.444 | 0.238 | 0.367 | 0.396 | 0.524 |
| E1 600M | Sequence Only | 0.475 | 0.482 | 0.472 | 0.254 | 0.342 | 0.419 | 0.523 |
| Inference with Homologous Sequences / MSA in-context |  |  |  |  |  |  |  |  |
| PoET | Sequence + Homologs | 0.482 | 0.541 | 0.464 | 0.491 | 0.478 | 0.478 | 0.510 |
| E1 150M | Sequence + Homologs | 0.482 | 0.527 | 0.476 | 0.494 | 0.476 | 0.477 | 0.515 |
| E1 300M | Sequence + Homologs | 0.485 | 0.534 | 0.478 | 0.490 | 0.471 | 0.480 | 0.520 |
| E1 600M | Sequence + Homologs | 0.487 | 0.537 | 0.488 | 0.500 | 0.478 | 0.485 | 0.525 |

### B. Unsupervised contact map prediction benefits from homologous sequences during inference

Unsu-pervised contact map prediction can be used as an efficient proxy to test whether the model has learned to encode information about the 3D structures of proteins during pre-training. In this section, we compare the performance of `E1` with publicly available models on the long-range contact prediction task for protein sequences from CAMEO [30, 31] and CASP15 [32] targets. We use the Categorical Jacobian approach [8] to assess the model’s internal knowledge of residue–residue contacts in an architecture-agnostic manner and report precision-at-L (the percentage of top-L predicted contacts that are correct). We define a residue pair as being in contact if their C*β*-C*β* distance is *<* 8Å, and we define long-range contact as contact between residues separated by at least 24 positions in sequence space.

We also evaluate whether the model can exploit additional information from homologous sequences during inference to improve contact-prediction performance. Homologs are sampled using the same procedure described in the previous section, with MSAs generated by ColabFold from the UniRef dataset. In contrast to the variant-effect prediction experiments, we do not ensemble over multiple prompts; instead, we fix the context length to 8192 and the maximum query similarity to 0.95 and use a single prompt for evaluation.

#### Results

We observe from Table 3 that `E1` models outperform the ESM family of models at all scales when tested in single-sequence mode. Moreover, we see consistent gains in performance when including homologous sequences during inference, indicating that the model is able to leverage in-context evolutionary information to identify putative 3D contacts in a protein. Finally, we provide some illustrative examples from the CAMEO dataset in Figure 3 where retrieval augmentation yields markedly improved contact-map predictions relative to single-sequence inference.

**Table 3.** Unsupervised contact map prediction performance as measured by Precision@L for long range contacts.

| Model Name | Model Inputs | Long-range Precision@L |  |
| --- | --- | --- | --- |
|  |  | CAMEO | CASP15 |
| Inference with query sequence only |  |  |  |
| ESM2-150M | Sequence Only | 0.348 | 0.272 |
| ESM2-650M | Sequence Only | 0.423 | 0.342 |
| ESM2-3B | Sequence Only | 0.434 | 0.339 |
| ESMC-300M | Sequence Only | 0.425 | 0.342 |
| E1 150M | Sequence Only | 0.466 | 0.387 |
| E1 300M | Sequence Only | 0.493 | 0.401 |
| E1 600M | Sequence Only | **0.512** | **0.425** |
| Inference with Homologous Sequences / MSA in-context |  |  |  |
| MSA Pairformer | Sequence + MSA | 0.489 | 0.428 |
| E1 150M | Sequence + Homologs | 0.510 | 0.406 |
| E1 300M | Sequence + Homologs | 0.526 | 0.415 |
| E1 600M | Sequence + Homologs | **0.541** | **0.436** |

**Figure 1.**
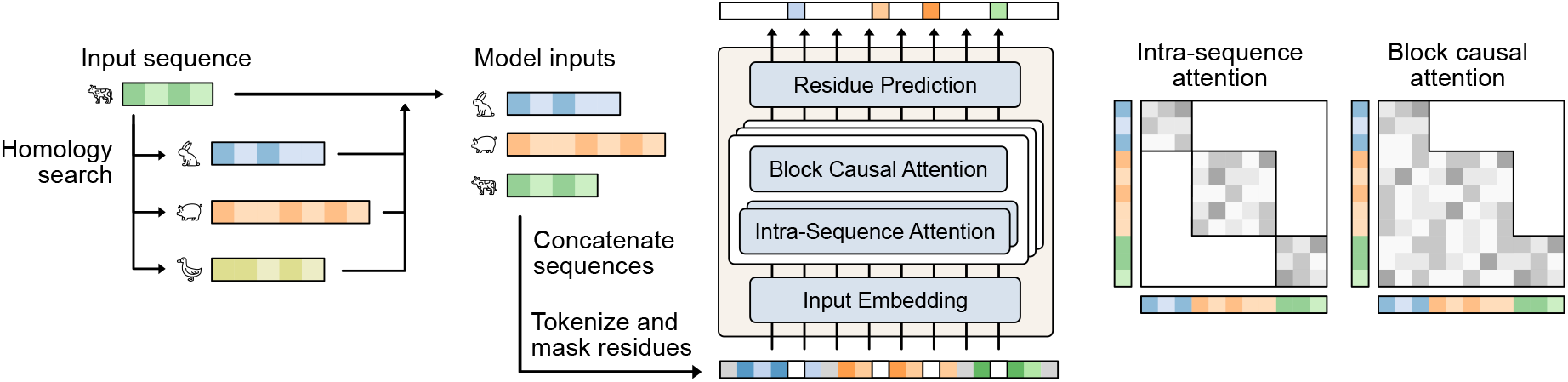
E1 Architecture. The `E1` model can take in homologous sequences in addition to an input query sequence. The homologous sequences are prepended to the query sequence to construct a multi-sequence input to the model. `E1` alternates between intra-sequence and block-causal attention, enabling it to build internal representations based on residues within the same protein sequence as well as residues in preceding homologous sequences within the concatenated multi-sequence input.

**Figure 2.**
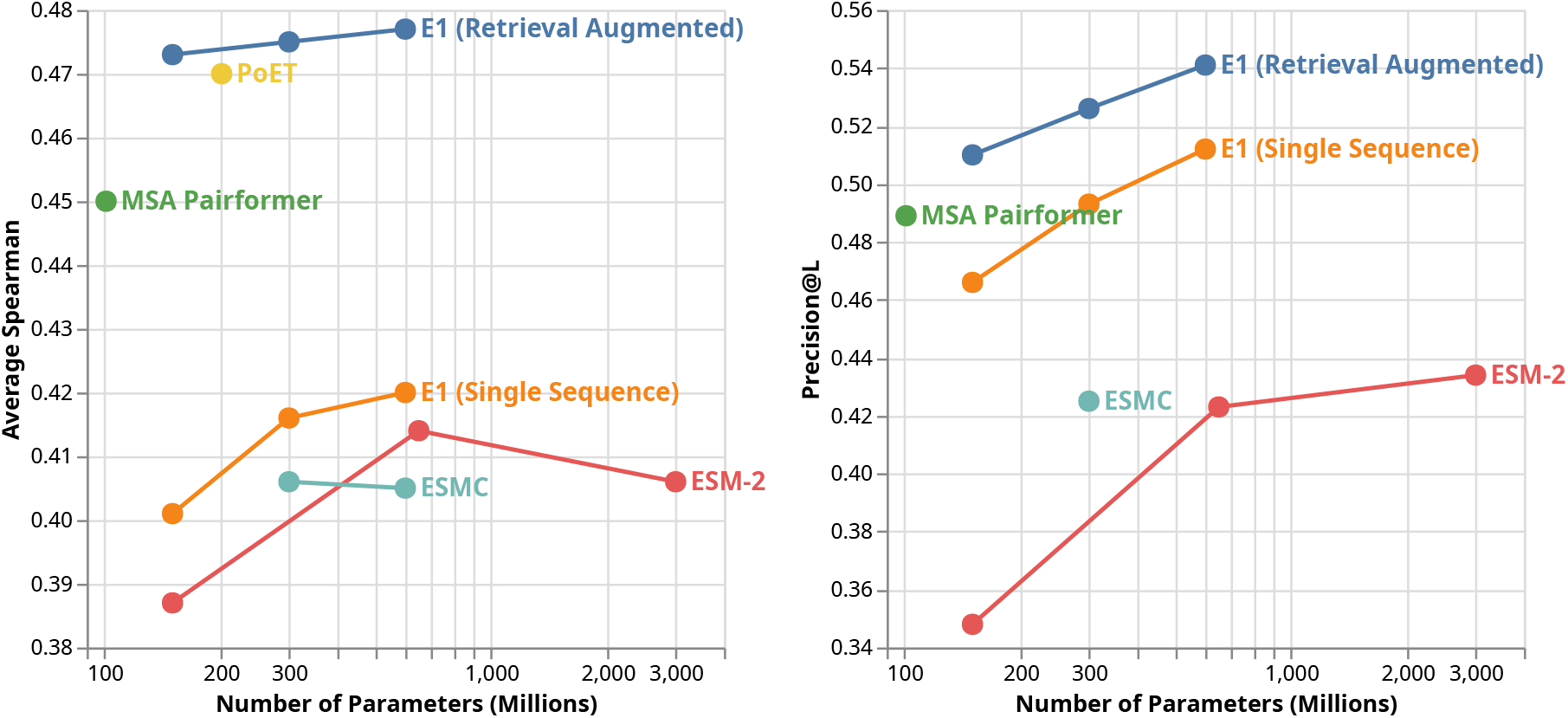
`E1` achieves state-of-the-art zero-shot performance compared to other publicly available PLMs in both sequence-only and retrieval-augmented mode. Scaling model parameters correlates with better performance. Left: Performance on Protein Gym substitution DMS Assays. Right: Unsupervised contact map prediction on a subset from CAMEO.

**Figure 3.**
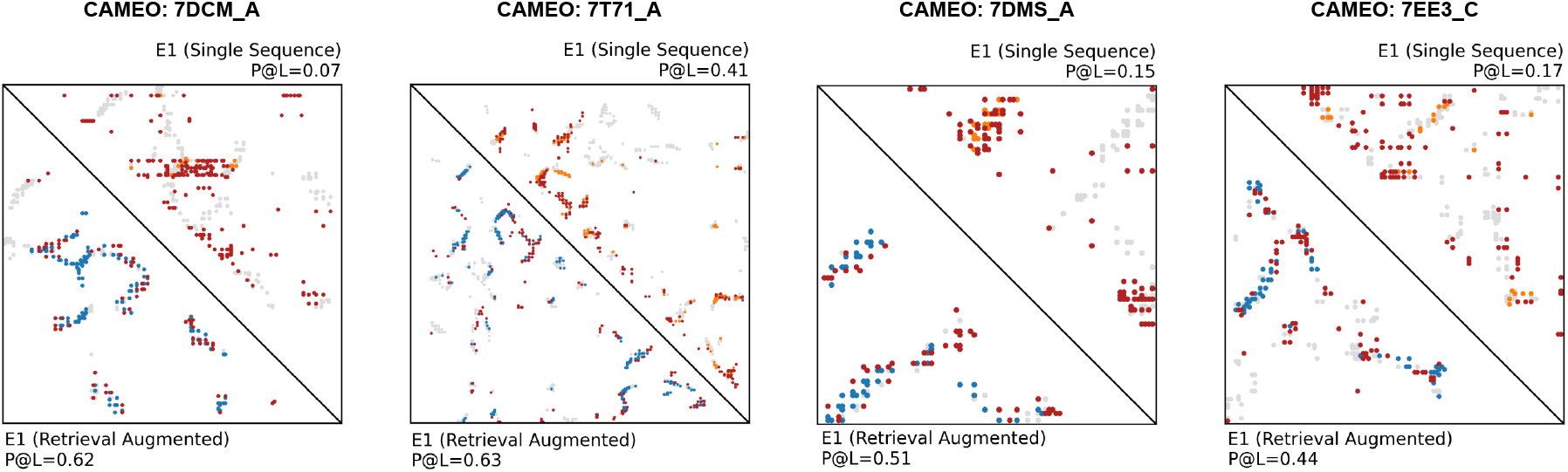
Examples from CAMEO dataset where retrieval augmentation helps `E1` identify contact it may have mispredicted when used in single sequence mode. Here, gray points are ground truth contacts, blue/orange points are correctly predicted contacts in retrieval-augmented/single-sequence mode, respectively, and red points are false positives.

## Discussion

We introduced **Profluent-E1**, a family of retrieval-augmented protein encoder models that can leverage unaligned evolutionarily related sequences at inference time to achieve superior performance. `E1` achieves state-of-the-art performance among publicly available models on variant-effect prediction (Protein Gym) and unsupervised contact-map prediction (CAMEO and CASP15), both in single-sequence mode and when augmented with homologs. We release three `E1` variants – 150M, 300M, and 600M – that are available for free for research and commercial use.

While we have shown the benefits of using retrieval augmentation on predictive performance for the `E1` family, several open questions remain regarding the inner workings of these models. In particular, further analysis is needed to disentangle how much `E1` is relying on the information encoded in the model weights during pre-training versus that derived from homologous sequences provided at inference time. Unlike other models like MSA Transformer [12], which may incorporate alignment information through specific attention mechanisms such as row and column only attention, `E1` models allow any residue in a given protein sequence to attend to any residue in preceding sequences within the multi-sequence input. This begs the question of whether the model implicitly learns to attend to positions that would have been aligned under a traditional MSA – or whether it exploits additional contextual signals from other regions of the homologous sequences beyond what alignment alone would provide.

Scaling laws seem to exist as we increase the model parameter count for our zero-shot evaluation tasks. However, we only extended this study to 600M parameters. Also, within the broader context of protein representation learning, we studied only sequence-based models to focus on the effects of retrieval augmentation. It has been shown that utilizing structural information in pretraining can lead to more efficient learning and more performant models in some contexts [16, 26, 33, 34]. Finally, it remains to be seen whether prompting the `E1` models with sequences that have specific properties can implicitly guide the model towards particular areas of the fitness landscape (for example, enzymes that work at specific pH levels or in specific organisms) and thereby optimize for desired functional attributes. We hope that by making these models publicly available under a permissive license, the research community will be able to provide answers to these and other questions, helping to develop more capable RA-PLMs in the future.

Overall, the Profluent-E1 family of models demonstrates the continued value of research in improving protein language models and provides a new foundational tool for AI-driven protein design that advances both predictive performance and practical utility for a large class of protein design workflows.

## Code availability

We make inference code and model weights available at https://github.com/Profluent-AI/E1 under a permissive license. See license details here: https://github.com/Profluent-AI/E1/blob/main/NOTICE

## Author contributions

**Data:** Sarthak Jain, Joel Beazer

**Pre-training:** Sarthak Jain, Aadyot Bhatnagar

**Evaluations:** Sarthak Jain, Jeffrey A. Ruffolo

**Overall Scientific Direction:** Sarthak Jain, Ali Madani

## Footnotes

∗ The metrics for MSA Pairformer are taken from the original paper, while PoET, ESM-2, and ESMC are sourced from the Protein Gym public leaderboard

